# The Open Constellation of Nutritional Values: Mapping the Health Star Rating System

**DOI:** 10.1101/2025.11.05.686692

**Authors:** Valentin Lafargue, Flore N’kam Suguem

## Abstract

**Background:** Nutritional awareness is increasing worldwide due to rising obesity rates and the widespread use of nutrition-tracking technologies. However, nutritional labels on food packaging remain difficult for many consumers to interpret. To improve clarity, public authorities have introduced simplified front-of-pack labeling systems, such as the Health Star Rating (HSR) used in Australia and New Zealand, which summarizes a product’s overall nutritional quality. Despite its adoption, the lack of open data on Health Star Rating labeled products limits independent research, reproducibility, and data-driven policy evaluation.

**Methods:** We introduce OpenHSR, the first open, FAIR-compliant dataset (Findable, Accessible, Interoperable, and Reusable) dedicated to the Health Star Rating system. The dataset includes 246 unique food products collected from Australian supermarket websites and brand databases. Using these data, we trained and compared several regression models linear, tree-based, and neural approaches to predict Health Star Rating values from nutritional attributes, thereby developing a transparent, interpretable (“white-box”) Health Star Rating calculator.

**Results:** Machine learning models achieved strong predictive performance, with the Support Vector Regression model yielding the best overall accuracy (Mean Squared Error = 0.28). The trained models were then applied to predict Health Star Rating values for 6,554 Open Food Facts products, demonstrating scalability and generalizability of the approach.

**Conclusions:** OpenHSR provides the first openly available, standardized dataset linking nutritional composition and Health Star Rating values. Its transparent design supports reproducible research, automated Health Star Rating estimation, and cross-dataset integration. This resource enables future studies in nutrition science, food labeling policy, and machine learning applications for public health.

## 1 Introduction

The growing public focus on nutrition and health has heightened the need for transparent, data-driven evaluations of food quality. While many countries have implemented front-of-pack labeling systems to guide consumers toward healthier choices, researchers and policymakers often lack open access to the data required to independently assess their effectiveness. The Health Star Rating (HSR) system, used in Australia and New Zealand, is one such front-of-pack scheme. Similar to other systems, such as the Nutri-Score widely used in European countries, HSR aims to support consumer decision-making by providing a clear, standardized indication of a product’s overall nutritional profile. These labels typically appear on packaged foods and range from 0.5 to 5 stars. Despite the broad adoption of HSR, access to large-scale, structured datasets of HSR-labeled products remains limited, restricting the potential for comprehensive research and policy evaluation.

Existing datasets such as Open Food Facts, FoodSwitch, and AUSNUT 2023 primarily focus on providing nutrient information, either as open data or through consumer-oriented applications that promote healthier food choices. However, a gap remains, as none of these datasets specifically emphasize or comprehensively capture HSR information.

To address this gap, we introduce OpenHSR, the first open and publicly available dataset dedicated to the Health Star Rating system. The dataset aggregates information from multiple Australian supermarkets and includes each product’s name, nutritional composition per 100 g or 100 mL, HSR category, and assigned Health Star Rating. Our dataset has been developed in accordance with the FAIR principles Findable, Accessible, Interoperable, and Reusable to ensure high-quality data management and research reproducibility. By making this dataset openly accessible, our goal is to enable reproducible analyses of nutritional trends, facilitate evaluation of the HSR system’s scoring consistency, and support further research in nutrition science, food policy, and machine learning applications for health-related data.

In Section 2, we describe the HSR system in detail. Section 3 introduces the datasets available for nutritional analysis, including publicly accessible ones for the Nutri-Score, but none for HSR. In Section 4, we present our dataset and its diversity in HSR coverage. Finally, Section 6 introduces machine learning–based regression algorithms to predict HSR from nutritional values, enabling the creation of our own HSR calculator and checking of the coherence between the actual calculation and the theoretical guidelines.

## 2 Health Star Rating System introduction

The Health Star Rating abbreviated HSR is the outcome which our dataset is focused on [9]. The Health Star Rating System, jointly implemented by the Australian and New Zealand governments, assigns packaged foods a rating from 0.5 to 5 stars. The higher the rating, healthier products. The HSR are computed through the official HSR Calculator, which applies the Nutrient Profiling Scoring Criterion (NPSC) to determine a product’s category-specific score and corresponding star rating. First, the products are classified into categories:

- 1. Non-dairy beverages, jellies and water-based ice confections
- 1D. Milk and Dairy beverages (and alternatives)
- 2. Foods
- 2D. Dairy foods (and alternatives)
- 3. Oils and Spreads
- 3D. Cheese

and then reclassified into two different categories:

1. Milk and Dairy beverages (and alternatives), foods including dairy foods (and alternatives), oils, spread and cheese.
2. Non-dairy beverages, jellies and water-based ice confections

For each product in the first category, the baseline Health Star Rating (HSR) is calculated using four negative components, as these are associated with increased risk factors for chronic diseases: energy, saturated fat, sodium, and total sugars. To obtain the final rating, the baseline score is then adjusted to account for the presence of positive components, namely protein content, amount of fruits, vegetables, nuts, and legumes (FVNL) and dietary fiber in the product, all of which contribute to improving the initial baseline rating. For products belonging to the second category, the HSR calculation follows a similar approach; however, among the components associated with the risk of chronic disease, only energy and total sugars are considered in determining the baseline score. The final Health Star Rating is then derived after incorporating the positive components, resulting in a single value that reflects the overall nutritional quality of the product.

## 3 Related Datasets

Open Food Facts is an open-source, publicly available dataset accessible at https://fr.openfoodfacts.org/ [12], containing metadata, Nutri-Score, NOVA classification, and Green Score for a wide variety of food products. The primary method of data collection relies on user submissions from around the world. While Health Star Rating (HSR) information is available for some products, there are often missing values in the metadata. Moreover, HSR data is not included in the downloadable version of the dataset, which limits its reusability for research purposes.

FoodSwitch is primarily a mobile phone application designed to help consumers make healthier food choices while enabling crowdsourcing of national food composition data [6]. Although the app includes Front-of-Pack nutrition labels such as the Health Star Rating (HSR), the underlying dataset is not directly accessible. Access requires submitting a request and demonstrating a research purpose, which limits its usability for broader projects. Consequently, the dataset does not fully comply with FAIR principles, as it is not readily findable, accessible, or reusable for a wide range of users.

The AUSNUT dataset is a comprehensive collection of detailed food supply data that supports the 2023 National Nutrition and Physical Activity Survey [8] and is accessible at https://www.foodstandards.gov.au/science-data/ food-nutrient-databases/ausnut. Classification or coding systems are employed to group similar foods for reporting intakes and tracking trends over time. The study utilized three classification schemes to report:

- Foods, dietary supplements, and nutrient intakes
- Intake of discretionary versus non-discretionary foods
- Nutrient intakes relative to the Australian Dietary Guidelines

However, this dataset does not include Health Star Rating (HSR) information, limiting its applicability for analyses that rely on front-of-pack nutritional labeling metrics.

## 4 Dataset Description

### 4.1 Overview

The OpenHSR dataset comprises 246 unique food products available in Australian retail chains in 2025 [11]. Each record contains approximately 20 fields, which can be grouped into three main categories:

Nutrient composition includes energy (kJ and kcal), protein, total fat, saturated fat, carbohydrate, sugars, fiber, and sodium. Metadata includes product name, brand, product type, category, serving size, package weight, country of origin, retailer, date collected, allergens, ingredients, and data source. Health Star Rating (HSR) is provided as a score from 0.5 to 5, reflecting overall nutritional quality.

### 4.2 Data Collection Process

The data was manually obtained from the package images of products from retailers websites and Brand-specific online pages such as Coles, Woolworths, Kellogg’s, Aldi and Fourntwenty. A part of the data was also similarly obtained from Open data facts’ website. Using the packaging information is known to be more reliable especially in the case of open data facts where the written information on their website differed from the information on the packaging in some instances, decreasing the risk of human-induced errors.

Manual collection ensured accuracy and consistency in variable representation, while metadata fields were harmonized to allow cross-product comparisons.

### 4.3 Data Validation

The dataset is provided in a tabular format (CSV), where each row represents a product and each column corresponds to a distinct variable. The dataset has no missing values in numerical entries.

Both manual and computational double-checks of nutritional panels and Health Star Ratings (HSR) were conducted to ensure a high-quality dataset.

The dataset also integrates the FAIR principles to support reproducibility and facilitate future research use:

#### Findable

Each product record is uniquely identified, and the dataset is assigned a DOI via Open Research Europe.

#### Accessible

The dataset is openly available under a CC BY 4.0 license, with direct download access.

#### Interoperable

Standardized units, consistent variable names, and harmonized categories enable integration with other datasets and computational pipelines.

#### Reusable

Comprehensive metadata, data dictionaries, and clear documentation support reproducibility, machine learning applications, and policy evaluations.

We described every variable gathered in the OpenHSR dataset in our GitHub.

## 5 Exploratory Analysis

The dataset exhibits sufficient diversity in HSR, which has enabled the development of a transparent, interpretable white-box HSR calculator.

The distribution of product categories across HSR reveals how nutritional quality varies within and between categories, as observed in Fig. 1a. Some categories tend to achieve higher HSRs like the 1D category, while other like the 2D category tend to have lower ratings, reflecting differences in their nutrient profiles. This analysis highlights the relationship between product type and nutritional quality.

**Fig. 1:**
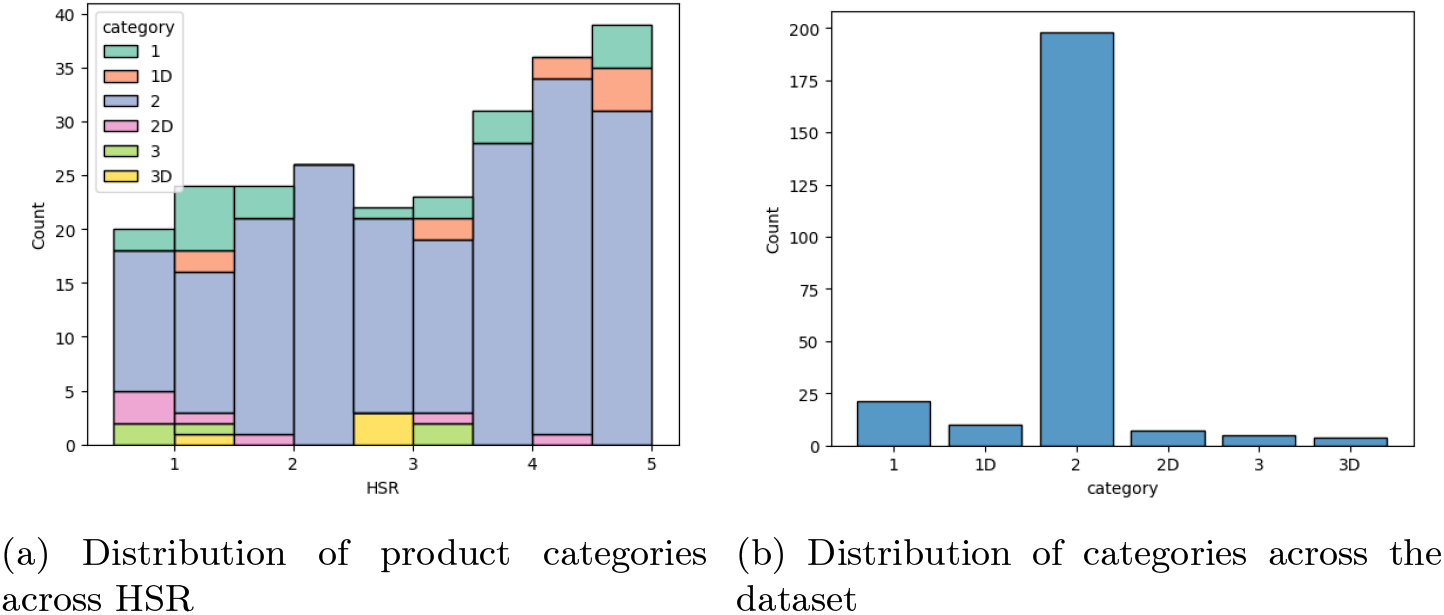
Descriptive statistics of the OpenHSR

Fig. 1b shows the categories’ distribution in the dataset, it provides insight into its overall composition. The distribution of the different categories are heterogeneous; however, this is not a characteristic of our dataset but the general distribution in supermarket. Indeed, we do not expect the same number of product in the ‘2’ category being `Food` as in the ‘3D’ category, which is `Cheese`.

We refer to Fig. 2a for the nutritional values’ distributions across the dataset, they confirm the variability among the recorded food items. Moreover, a clear relationship is observed between energy content and HSR, products with higher calories per 100g tend to have lower HSRs, and conversely, products with lower calorie content tend to have higher HSRs. In Fig. 2b, lighter-colored points represent lower HSRs, which occur less frequently but correspond to higher energy content per 100g.

**Fig. 2:**
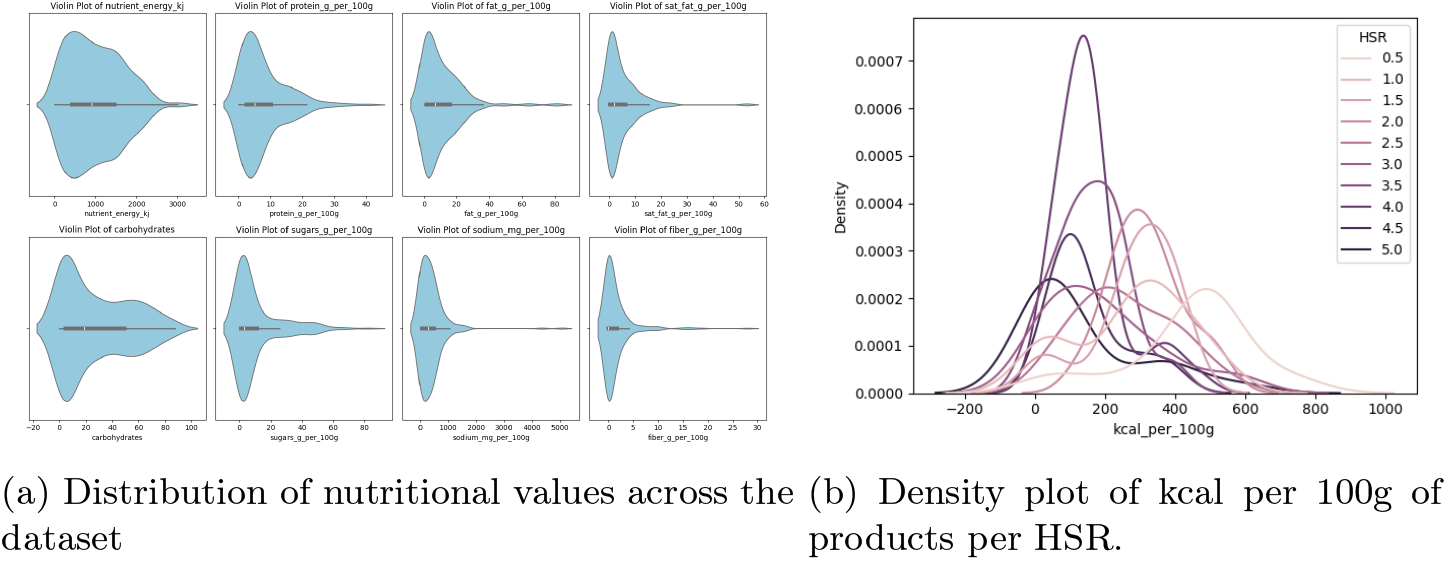
Descriptive statistics of the nutritional values within the OpenHSR

## 6 Health Star Rating Calculator: Benchmark

We used a diverse set of machine learning (ML) regression models to predict the Health Star Rating (HSR). As summarized in Table 1, the models include three fully white-box and interpretable algorithms (LR, DT, kNN), two ensemble methods (RF, XGBoost)(whose individual base learners are interpretable although the overall ensemble is not), and three black-box models (SVR, MLP, FTN). A detailed description of our model implementation and deployment strategy is provided in our GitHub.

**Table 1:** Performance of various machine learning regression algorithms for HSR prediction. Each algorithm was evaluated based on the Mean Squared Error (MSE), accuracy (acc) after rounding regression outputs to the nearest 0.5-star increment, and lenient accuracy (by-half-acc), which tolerates a 0.5-star deviation.

| ML regression algorithm | MSE | acc | by-half-acc | Interpretable |
| --- | --- | --- | --- | --- |
| Linear [10] | 0.82 | 0.42 | 0.76 | ✓ |
| Decision Tree [2] | 0.55 | 0.34 | 0.76 | ✓ |
| K-Nearest Neighbor [7] | 0.40 | 0.42 | 0.86 | ✓ |
| Random Forest [1] | 0.39 | 0.32 | 0.84 | — |
| XGBoost [3] | 0.35 | 0.54 | **0.92** | — |
| Support Vector Machine [5] | **0.28** | 0.62 | 0.86 | ✗ |
| Multilayer Perceptron [13] | 0.67 | 0.40 | 0.78 | ✗ |
| Fine-tuned network [13] | 0.86 | 0.26 | 0.50 | ✗ |

Overall, the models achieved strong predictive performance for HSR. For example, XGBoost achieved a half-accuracy of 92% with a mean squared error below 0.4. In contrast, fully interpretable models such as the Decision Tree (see Fig. 3) produced lower predictive accuracy compared to less interpretable approaches, notably the Support Vector Machine (SVM) regression, which was among the best performing models.

**Fig. 3:**
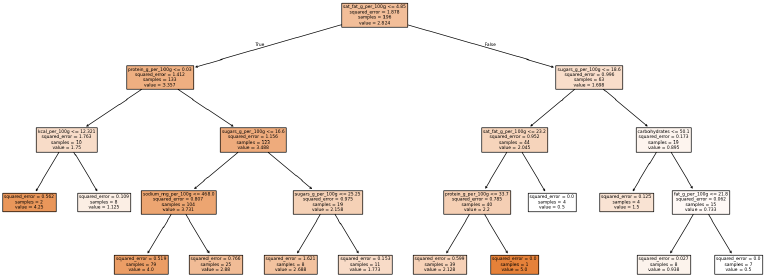
Decision-making process of the decision tree Regression algorithm. It shows which variable it uses, and the HSR predicted in consequences. The lighter-colored the cells are, the lower the HSR predicted will be.

Performance could likely be improved, particularly for models such as SVM and Multilayer Perceptron (MLP), with access to additional labeled data. To address data limitations, we attempted to pre-train a foundation model to predict Nutri-Scores, subsequently using its latent representations to predict HSR. However, this approach did not yield satisfactory results for HSR prediction (Table 1), despite performing well for Nutri-Score prediction (MSE = 0.24, after transforming Nutri-Scores to a 1–5 numerical scale).

From an applied perspective, although a half-star difference in HSR may not substantially alter consumer judgments, prioritizing interpretability over marginal performance gains can be justified. However, for research applications or large-scale predictive analyses, the SVM regression model is recommended due to its superior performance (lowest MSE). To demonstrate the generalizability of our approach, Section 7 presents HSR predictions on an independent nutritional dataset.

## 7 Health Star Rating Predictions

We applied our trained models to the OpenFoodFacts dataset by selecting all products containing the nutritional attributes required by the algorithms described in the previous section. The only missing feature was the category variable introduced in Section 2. To address this, we constructed a mapping dictionary linking the most common categories in our training data to their corresponding categories in the HSR classification system.

Using the Support Vector Regression (SVR) model, we then predicted HSR values for the selected products. Although direct validation of these predicted scores was not possible due to the absence of ground-truth HSR labels, we performed a consistency check by examining the resulting distribution of predicted ratings. As shown in Fig. 4, the distribution appears well diversified, supporting the reliability and generalizability of our HSR estimation approach. Notably, our method demonstrated strong computational efficiency, producing HSR predictions for 6,554 products in only 0.1 seconds, suggesting its suitability for large-scale applications.

**Fig. 4:**
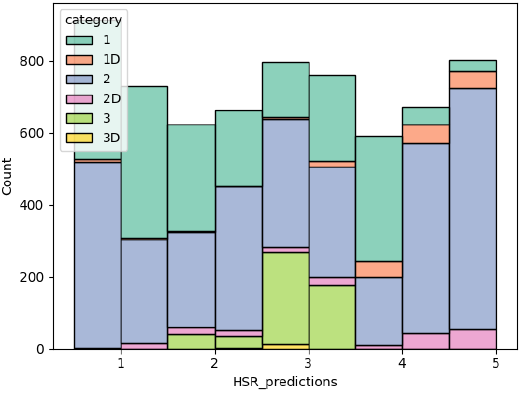
Distribution of product categories across HSR for the sampled OpenFoodFacts dataset using SVR

## 8 Conclusion & Perspective

In this work, we presented the creation of a new dataset linking Health Star Ratings (HSR) with detailed product metadata for Australian retail foods. We provided a thorough analysis of the dataset, including its structure, data quality, and potential applications. Additionally, we demonstrated how machine learning models trained on this dataset can be used to predict HSR values for products where this information is not available, highlighting its utility for research and consumer applications.

The dataset is publicly accessible online, and its design adheres to the FAIR principles, ensuring it is findable, accessible, interoperable, and reusable. We anticipate that this resource will facilitate further studies on nutrition labeling, dietary assessment, and predictive modeling in food and health research.

### Limitations

One of the main limitations of our dataset is its relatively small size (246 entries), which may affect the generalizability and performance of machine learning models. In addition, the dataset focuses exclusively on products available in the Australian market, limiting its applicability to other regions. Due to the fact that the data were collected manually, it may be subject to packaging updates and labeling changes over time. Furthermore, serving-size variability is not included, which may influence nutrient comparisons across products.

### Future Directions

To address these limitations, we plan to expand the dataset by increasing the number of product entries and incorporating data from multiple regions. Future versions will aim to automate data collection processes, include serving-size information, and ensure periodic updates to maintain accuracy and relevance.

## Ethics and consent

Ethical approval and consent were not required.

## Data availability

### Underlying data

Zenodo: *Open Health Star Rating (OpenHSR) Dataset*. https://doi.org/10.5281/zenodo.17469191

Data are available under the terms of the Creative Commons Attribution 4.0 International license (CC BY 4.0).

### Software availability

Source code available at: ValentinLafargue/HealthStarDataset The GitHub repository is organized this way:

1. Data folder: OpenHSR.csv is our original data file explained above, Open-FoodFacts_clean_sample.csv is a dataset sample from OpenFoodFacts which we create by selecting non-missing values on variable of interest and mapping the former categories to HSR categories.
2. NN folder: We kept the neural network’s weights for reproductibility.
3. HS_description.ipynb: Notebook which describes the dataset and enabled us to create the graphs presented in our paper.
4. HS_prediction.ipynb: Notebook used to produce the paper benchmark, it predicts the HSR using classical machine learning and neural networks.
5. requirements.txt: librairies needed in the project
6. utils.py: python fonctions used in the neural network

The software is released under the Creative Commons Attribution 4.0 International license (CC BY 4.0).

## Declaration of generative AI and AI-assisted technologies in the writing process

During the preparation of this work the authors used ChatGPT in order to enhance the readability and language. After using the tool, the authors reviewed and edited the content as needed and take full responsibility for the content of the publication.

## References

1. Breiman, L.: Random forests. Machine Learning 45, 5–32 (2001). 10.1023/A:1010933404324, https://www.stat.berkeley.edu/~breiman/randomforest2001.pdf

2. Breiman, L., Friedman, J.H., Olshen, R.A., Stone, C.J.: Classification and Regression Trees. Wadsworth International Group, Belmont, CA (1984), https://www.routledge.com/Classification-and-Regression-Trees/Breiman-Friedman-Stone-Olshen/p/book/9780412048418

3. Chen, T., Guestrin, C.: Xgboost: A scalable tree boosting system. In: Proceedings of the 22nd ACM SIGKDD International Conference on Knowledge Discovery and Data Mining (KDD ‘16). pp. 785–794 (2016). 10.1145/2939672.2939785, https://arxiv.org/abs/1603.02754

4. Defazio, A., Yang, X., Mehta, H., Mishchenko, K., Khaled, A., Cutkosky, A.: The road less scheduled (2024)

5. Drucker, H., Burges, C.J.C., Kaufman, L., Smola, A.J., Vapnik, V.N.: Support vector regression machines. In: Advances in Neural Information Processing Systems 9 (NeurIPS 1996). pp. 155–161 (1997), https://papers.neurips.cc/paper/1238-support-vector-regression-machines.pdf

6. Dunford, E., Trevena, H., Goodsell, C., Ng, K.H., Webster, J., Millis, A., Goldstein, S., Hugueniot, O., Neal, B.: Foodswitch: A mobile phone app to enable consumers to make healthier food choices and crowdsourcing of national food composition data. JMIR mHealth and uHealth 2(3), e37 (2014). 10.2196/mhealth.3230

7. Fix, E., Jr., J.L.H.: Discriminatory analysis: Nonparametric discrimination – consistency properties. Tech. Rep. 4, USAF School of Aviation Medicine (1951), https://cs.nyu.edu/home/people/in_memoriam/roweis/csc2515-2006/readings/fix_hodges_51.pdf

8. (FSANZ), F.S.A.N.Z.: Ausnut 2023 – the 2023 national nutrition and physical activity study food and nutrient database. https://www.foodstandards.gov.au/science-data/food-nutrient-databases/ausnut (2025), accessed: 2025-10-30

9. Health Star Rating Unit, N.P.S.: Hsr system calculator and style guide (2025)

10. Legendre, A.M.: Nouvelles méthodes pour la détermination des orbites des comètes. Courcier, Paris (1805), https://archive.org/details/nouvellesmthode00legegoog

11. N’kam Suguem, F., Lafargue, V.: Open health star rating (openhsr) (version v1). Data set (2025). 10.5281/zenodo.17469191, https://doi.org/10.5281/zenodo.17469191

12. Open Food Facts Team: Open food facts – a collaborative, open database of food products from around the world. https://data.europa.eu/en/publications/use-cases/open-food-facts-collaborative-open-database-food-products-around-world (2018), accessed: 2025-10-30

13. Schmidhuber, J.: Deep learning in neural networks: An overview. Neural Networks 61, 85–117 (Jan 2015). 10.1016/j.neunet.2014.09.003, http://dx.doi.org/10.1016/j.neunet.2014.09.003

